# Coordinate Based Random Effect Size meta-analysis of neuroimaging studies

**DOI:** 10.1101/089565

**Authors:** CR Tench, Radu Tanasescu, WJ Cottam, CS Constantinescu, DP Auer

## Abstract

Low power in neuroimaging studies can make them difficult to interpret, and Coordinate based meta‐analysis (CBMA) may go some way to mitigating this issue. CBMA has been used in many analyses to detect where published functional MRI or voxel-based morphometry studies testing similar hypotheses report significant summary results (coordinates) consistently. Only the reported coordinates and possibly *t* statistics are analysed, and statistical significance of clusters is determined by coordinate density.

Here a method of performing coordinate based random effect size meta-analysis and meta-regression is introduced. The algorithm (ClusterZ) analyses both coordinates and reported *t* statistic or *Z* score, standardised by the number of subjects. Statistical significance is determined not by coordinate density, but by a random effects meta-analyses of reported effects performed cluster-wise using standard statistical methods and taking account of censoring inherent in the published summary results. Type 1 error control is achieved using the false cluster discovery rate (FCDR), which is based on the false discovery rate. This controls both the family wise error rate under the null hypothesis that coordinates are randomly drawn from a standard stereotaxic space, and the proportion of significant clusters that are expected under the null. Such control is vital to avoid propagating and even amplifying the very issues motivating the meta-analysis in the first place. ClusterZ is demonstrated on both numerically simulated data and on real data from reports of grey matter loss in multiple sclerosis (MS) and syndromes suggestive of MS, and of painful stimulus in healthy controls. The software implementation is available to download and use freely.

## 2 Introduction

Neuroimaging studies often involve few subjects and have low statistical power to detect true effects, and with lack of power comes increased risk that significant results are false positives (Button 2013). Add to this the common use of uncorrected p-value thresholds (Bennett 2009), and neuroimaging studies can become difficult to interpret. This situation may be compounded if the data violate the methodological assumptions of the analysis (Eklund 2016). Meta-analysis can be used to synthesize the evidence across similar neuroimaging studies going some way to mitigating these problems (Ioannidis 2005), and there are various methods of statistically combining the results (Lazar 2002). Image based meta-analysis (IBMA) is the most powerful approach, but is currently limited by availability of suitable statistical images. Coordinate based meta-analysis (CBMA), on the other hand, uses just the available summary reports (coordinates and possibly *Z* scores or *t* statistics) from functional MRI/PET or voxel‐based morphometry studies measuring common effects, and has been utilised in many published studies; the aim is similar to that of IBMA within the limits of the available data (Salimi-Khorshidi 2009). The results of CBMA consist of clusters of coordinates where studies have reported significant effect in similar anatomical locations, representing consistency and indicating relevancy of brain structures, while coordinates not recruited into clusters are considered study specific. Popular CBMA algorithms include the activation likelihood estimate (ALE) (Turkeltaub 2002, Laird 2005, Eickhoff 2009, Eickhoff 2012, Turkeltaub 2012) and the multi-level kernel density (MKDA) algorithm (Wager 2007). Signed differential mapping (SDM) (Radua 2010) is similar to the ALE but incorporating the sign of effect at the reported coordinates to distinguish grey matter loss from grey matter increase, or fMRI activation from deactivation. Effect size SDM (ES-SDM) (Radua 2012) takes this further and uses the reported *t* statistic associated with each coordinate, and can also incorporate statistical parametric maps.

There are technical issues with these CBMA algorithms that limit interpretability and specificity. Firstly statistical tests are performed voxel-wise making the relevant cluster-wise type 1 error rates difficult, if not impossible, to assess. Secondly the significance is, at least in part, determined by the density of coordinates from different experiments meaning that coordinates forming a small cluster are more significant than if they formed a larger cluster. Yet it is not clear, for example, that studies reporting thalamic coordinates producing a cluster over the thalamic volume should be less significant than the same studies reporting coordinates producing a smaller cluster in the smaller putamen structure. Finally, the uncorrected p-value threshold employed by both SDM and ES-SDM does not control the type 1 error rate in a principled way (Bennett 2009), and without estimated error rates there is no way to assess the significance of the results given the ~2×10^5^ voxel-wise statistical tests; this may propagate the very problems the MA was employed to mitigate (see (Tanasescu 2016) for example).

LocalALE is a CBMA algorithm (Tench 2013, Tench 2014) that addresses some of these limitations. It employs an interpretable cluster-level type 1 error rate control scheme, the false cluster discovery rate (FCDR), made possible by performing statistical tests at the coordinate, rather than the voxel, level. The results are such that at-most some specified proportion of the clusters declared significant are expected under the null hypothesis. LocalALE also adjusts its parameters to avoid false negatives when there are few studies and avoid false positives when there are many studies. Furthermore, LocalALE assigns coordinates to clusters in a binary fashion (belonging to a specific cluster, or no cluster), and as a consequence can analyse positive and negative effects (activation and deactivation, for example) simultaneously, allowing post-hoc checks for sign consistency. Nevertheless, LocalALE is unable to utilise the sign or magnitude of the reported effect to perform statistical inference, and the cluster significance is determined by coordinate density biasing the results to smaller clusters.

Here a new coordinate based random effect size (CBRES) meta-analysis (MA), and meta-regression, method (ClusterZ) is detailed. The algorithm deviates from other CBMA methods by performing inference on a standardised effect size, which is related to the Z score or *t* statistic reported by most studies. Consequently the density of coordinates within cluster does not influence statistical significance, so large and small clusters are considered on an equal footing. A random effects meta‐analysis approach is taken and model parameters are estimated by maximum likelihood estimation (MLE) and significance assessed by comparing models using a likelihood ratio test (LRT). Models can be devised to test for evidence of a non-zero effect size, effect size difference between groups, or significant linear regression. ClusterZ also requires consistent spatial effect across studies for significance, and uses this to control the type 1 error rate such that quantifiably more clusters are declared significant than are expected if the studies report uncorrelated spatial effects. Furthermore, it adjusts parameters to avoid false positive and false negative results depending on the number of studies. ClusterZ is closer to traditional MA in that estimates of effect and variance are computed. It provides an alternative to ES-SDM for coordinate based meta-analysis but with the advantage that the type 1 error rate is controlled, quantified, and interpretable.

## 3 Methods

There are several steps to the ClusterZ algorithm, detailed below. In summary, clusters are formed by reported coordinates that are more densely packed than average. Then, a random effects analysis is performed to give a p-value in each cluster. The same analysis is then performed on many pseudo experiments, in which each coordinate has been replaced by a random one to simulate studies reporting spatially uncorrelated effects. Declaration of significance in ClusterZ has two requirements: 1) that within cluster there is a consistent effect size reported such that the p-value is small, and 2) for a given p-value threshold the number of observed clusters with smaller p-values is quantifiably greater than average for the pseudo experiments. The second requirement indicates how ClusterZ controls the type 1 errors through the false cluster discovery rate.

### 3.1 Cluster forming

The clustering algorithm is identical to that used by LocalALE, and is detailed in (Tench 2013) but recapped here. It is based on a popular algorithm: density based spatial clustering of applications with noise (DBSCAN) (Ester 1996). The aim is to produce clusters of densely packed coordinates while not recruiting coordinates outside these clusters, which DBSCAN considers noise; in the present application these coordinates are considered study specific effects rather than noise. The initial step is a measure of overlap of coordinates in different studies. A coordinate that overlaps (they are separated by a distance <Δ) coordinates in *n* other studies has an overlap score of *n*. For a coordinate to be considered part of a cluster, its overlap score must be at least 3 according to the DBSCAN algorithm, since an overlap score of 2 or less means the coordinate is link in a chain, rather than a cluster, of coordinates. The peak of any cluster is the coordinate, or collection of coordinates, with the highest overlap score. The clustering algorithm proceeds by finding the peak coordinate that is not already assigned to a cluster and assigns it a cluster number. Coordinates that overlap members of this cluster, and have equal or lower overlap score, are recruited to the cluster. This continues until there are no more valid overlapping coordinates to be added to the cluster. The process then continues starting with the coordinate with the highest overlap score that is not already part of a cluster. The result is a set of clusters of coordinates that have a reducing (but not strictly) overlap score moving away from the peak.

The clustering process depends on the clustering distance Δ, which is analogous to the FWHM parameter used in other CBMA algorithms (Turkeltaub 2002, Radua 2010), and the algorithm to compute this has been detailed previously (Tench 2014). Briefly, however, the coordinates are redistributed randomly (see below) within an anatomical mask, which depending on the problem might be a grey-matter, white-matter, or whole-brain mask. For these coordinates a small value of Δ results in few coordinates having non-zero overlap scores, but this increases for larger Δ. It is helpful to consider the proportion of coordinates with non-zero overlap scores (divided by 2 to avoid coordinate A overlapping coordinate B being considered a second time as B overlapping A) as a function of the clustering distance: ϕ(Δ), the overlap fraction. The clustering distance used is that Δ which, on average, just causes each random coordinate to overlap with another in one other study such that ϕ(Δ)=0.5. With this value of Δ the clusters are formed from coordinates packed more densely than average for random coordinates, while the study specific coordinates that have lower than average density are unlikely to form clusters; for larger Δ the study specific coordinates will (wrongly) begin to form clusters, while for smaller Δ the true clusters may not form at all. This method adapts Δ to the number of studies, such that it is small for many studies because the density of coordinates is higher. Consequently the false negatives are reduced when there are fewer studies (the low density of coordinates when few studies are included requires Δ to be large so that clusters still form), while false positives are reduced when there are many studies (because the small Δ prevents the study specific coordinates being recruited into clusters) (Tench 2014); this is in contrast to the fixed FWHM parameter used in other algorithms that can paradoxically increase the false positives with an increasing number of studies (Tench 2014).

A subtlety in this algorithm is that independence of the coordinates within study cannot be guaranteed (Wager 2007, Eickhoff 2009). Reported coordinates that are very close to each other may be related, and the aim is to preserve this relationship. Independently randomising the coordinates within a mask using a uniform distribution would violate this aim, and therefore a more sophisticated approach is taken and has been described in detail previously (Tench 2013). Within-study clusters of coordinates (each coordinate separated by a distance <Δ) from at least one other coordinate from the same cluster) are formed. Each within-study cluster has a centroid, and the mean and standard deviation of the distances from the centroid, of the coordinates belonging to the cluster, computed. Each centroid is then randomly placed, with uniform probability, within the mask, and the cluster coordinates randomly placed about the centroid with the computed mean and standard deviation distance. A randomisation is rejected, and subsequently repeated until successful, if the within-study clusters of coordinates are randomised such that they overlap; see (Tench 2013) for details.

### 3.2 Models of effect size

In fMRI a biologically meaningful measure of effect is not available, although %BOLD (blood oxygenation level dependent) signal change has been suggested (Chen 2016). In VBM studies a more meaningful measure in terms of volume reduction is possible, but not routinely reported. Typically neuroimaging studies only report effect sizes reflecting statistical significance: the *t* score or Z score. These are dimensionless and relative effect sizes similar to Cohen’s *d* if scaled (Maumet 2016); *t* and Z are both scaled by subject numbers. The meaning, from a biological perspective, of such effects is not directly apparent, and often cannot be inferred from the report (Chen 2016). Prospective studies of regions known to be consistently reported as significant could be designed to answer this question, but first the regions must be identified and a statistical effect size estimated so that a sample size might be computed (Maumet 2016). ClusterZ can be used to provide this detail.

Within clusters the *Z* scores or *t* statistics are be combined to test if there is consistent statistical significance; publication bias or outlying effects, for example, might be reported by only one study and would not be consistent. Methods combining p-values (or equivalent), reviewed in (Lazar 2002), are appropriate for IBMA, where the test is for significant effect within a voxel given multiple statistical images. In CBMA, however, each reported coordinate has already been declared as significant by the reporting study, and the relevant question is about significant consistency of the reported effects across studies. Combining p-values (known to be significant already) is then not appropriate, so a meta‐analytic method of combining results from different studies (Lazar 2002) is employed instead, necessitating an effect size and estimate of variance.

Under certain assumptions the *t* statistic or *Z* score can be transformed and used as a dimensionless effect for MA or meta-regression. Assuming that studies report analyses of one or two groups of subjects then the *t* statistic, which has a student’s *t* distribution, is a measure of effect, or difference in effect between groups, reported in multiples of the standard error. The first difficulty with the *t* statistic as an effect size is that the standard error depends on the number of subjects, so larger studies appear to report larger effects. Therefore, the effect used in CBRES analysis is the *t* statistic, or *Z* score, scaled to obtain an estimate measured in multiples of the sample standard deviation, which does not depend on the number of subjects. A basic analysis is assumed, whereby simple *t* tests are performed. Then the effect size is computed from the *t* statistic by dividing by a function of the number of subjects *n*^*^

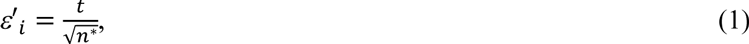

and the within-study variance is

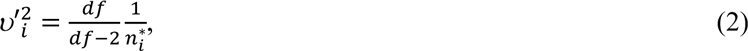

which includes the variance of the Student’s *t* distribution with *df* degrees of freedom. For a one sample study, for example just healthy control (HC) subjects, then *n^*^* is the number of subjects and *df=n^*^*−1. For a two sample study, such as a patient versus HC study, then

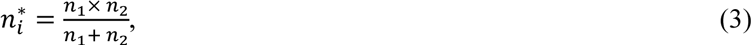

where *n*^1^ and *n*^2^ are the numbers of subjects in each sample and *df*=*n*_1_+*n*_2_-2; this assumes equal variance of effect in both groups. These standardised effect size and distributional assumptions assume a random effects approach has been used for analysis.

Under the given assumptions behind equations (1) to (3), they provide an effect size that is standardised against the number of subjects and have an associated variance estimate as required for MA. However, many studies employ a more sophisticated analysis and include multiple regressors (Friston 1994). This is partly corrected for by modifying the degrees of freedom of the *t* distributions by the number of extra regressors included; if the number of regressors is unknown, some underestimation of the effect variance will result. Furthermore, scaling the *t* statistic by 1/√n^*^ will not standardise to the same sample standard deviation across studies. This is not correctable, due to the limitations of reported effect, and will result in some between-study variance in effect size due to the range of analyses employed. Nevertheless, standardising the reported statistics should result in an effect with approximately known variance, and which tends to be larger for larger effects. Furthermore, ClusterZ attempts to make allowance for this limitation by including a between-study variance as a random effect.

One more assumption is made, for convenience, in what follows: that the subject numbers are sufficiently large that the reported *t* statistics are well approximated by Z scores such that the effect is

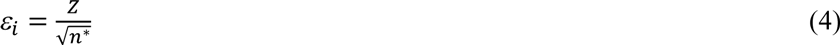

with variance

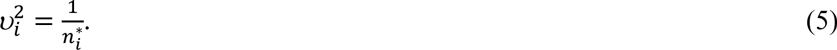

The *Z* scores are standardised by comparison to the range of possible Student’s *t* distributions making them easier to consider for the purpose of experimentation. However, ClusterZ does have the option to deal with either *t* of *Z* for effect sizes using either equations (1) and (2) or equations (4) and (5).

### 3.3 Random effect model

A random effects approach is employed to combine the effects given by equations (1-5), making some allowance for the often appreciable differences in experimental design and analysis between studies. To perform inference a random effect distribution must be assumed. Typically a normal distribution is assumed (Lazar 2002) so that for study *i*

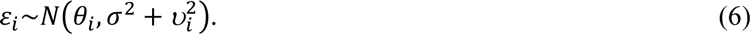

The parameter *θ_i_* is modified to reflect different models, and are estimated using MLE. The total variance is composed of the within-study variance 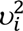 (equation (5)) and the between-study variance *σ*^2^, which is also estimated by MLE. With this model estimates are weighted more heavily to the larger studies due to the smaller within-study variance.

Various models are possible using different parameterisations of equation (6). To model the effect sizes by a grand mean the parameterisation is *θ_i_*=*μ*. For meta-regression the grand mean is modulated by a study specific covariate so *θ_i_*=*μ*−*βc_i_*, where *β* models the change in the grand mean due to the covariate *c*; this could be a continuous variable such as age, or a group indicator to investigate differences in effect size between groups.

#### 3.3.1 Maximum likelihood estimation with censoring

The maximum likelihood estimates of parameters *θ* and *σ* are generally straight forward to compute. However, a subtlety is that not all studies will report a coordinate and effect in every cluster, and those that don’t are censored by the statistical threshold used; for example a study applying an uncorrected p‐value threshold of 0·0001 reports only *Z* scores exceeding 3·72 in magnitude. Left, right, and interval censoring need to be considered. The log likelihood, which is maximised for parameter estimation, is a sum of contributions from uncensored and censored terms.

For uncensored data the log likelihood contribution for a single study is given by the probability (density) that effect size E=*ε_i_* given the parameters *θ_i_* and *σ* (P(E=*ε_i_*∣*θ_i_*,*σ*))

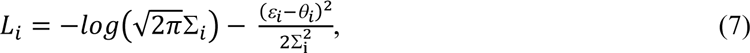

where

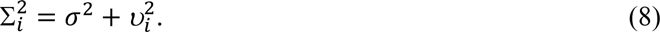

Left censoring occurs when a study reports a significant negative effect (for example deactivation) within a cluster but no effect size is given. If study *i* reports an effect size threshold such that E≤ −*T^i^*, where *T_i_* is the threshold magnitude, then the contribution to the likelihood is the probability P(E≤ − *T_i_*∣*θ^i^*,*σ*). Given the random effects model (equation (6)) this probability can be computed accurately using the error function (*erf*) (Chevillard 2012). The contribution to the log likelihood for a single left censored study is therefore

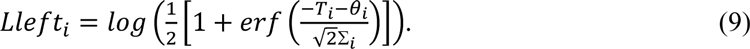

With right censoring study *i* reports an effect size threshold such that E≥ −*T^i^* and the contribution to the likelihood is the probability P(E≥ − *T_i_*∣*θ^i^*,*σ*), and log likelihood

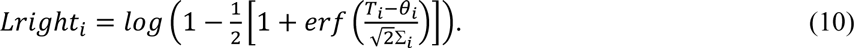

Interval censoring occurs when a study does not report a significant result within a cluster. In this case all that is known is that ∣E∣≤ −*T^i^*, and the likelihood is the probability P(∣E∣≤ − *T_i_*∣*θ^i^*,*σ*). In this case the contribution to the log likelihood is

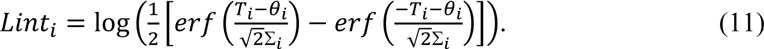

Summing these contributions over all studies gives the log likelihood

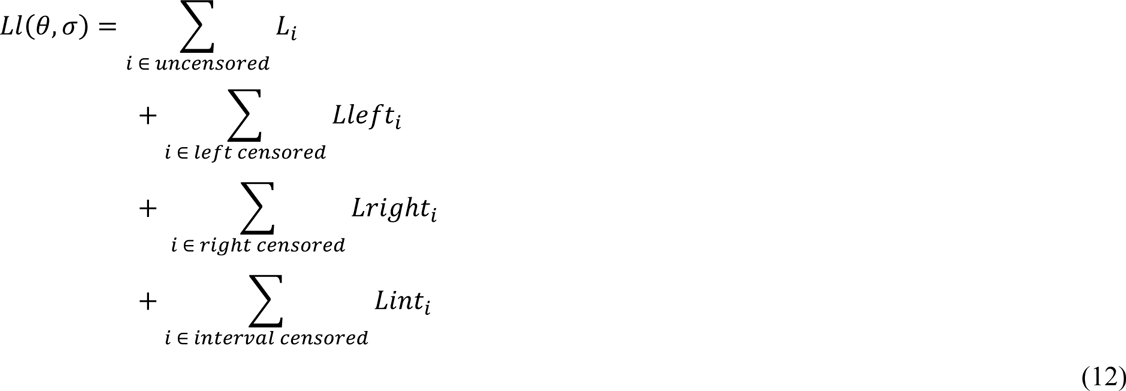

which is maximised for parameter estimation.

Effect size thresholds are often given in studies that employ uncorrected p-values. However, they are sometimes omitted. In the absence of a stated threshold, ClusterZ estimates them from the smallest magnitude effect size reported by the study. If no effect size is reported, then conservative low magnitude default threshold of 3.09 (corresponding to p<0.001) is used by default; a high default threshold might overestimate the effect and drive significance.

#### 3.3.2 Inference using the likelihood ratio test

Without the censoring, a one sample *t-*test or simple linear regression (SLR) model can be used for inference. However, neuroimaging study reports are censored so a likelihood ratio test, a general scheme for performing a statistical test by approximating the difference of two log likelihoods to a chi squared distribution, is used. The two likelihoods in question are the maximum likelihoods computed under the null and alternative hypotheses. To test the hypothesis that coordinates in a cluster have a non-zero grand mean effect size *μ* compute

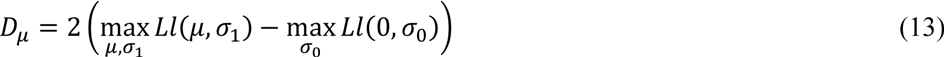

and compare *D_μ_* to a chi square distribution with one degree of freedom to compute the p-value. For the regression model, where *θ^i^*=*μ-βc_i_*, a test for non-zero regression coefficient *β* compares

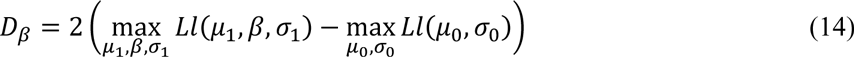

to a chi square distribution with one degree of freedom to compute the p-value.

### 3.4 Type 1 error control

The null hypothesis of the LRT’s is that there is zero mean effect. The null hypothesis used by ClusterZ and CBMA algorithms, however, is that studies report no common spatial effect. The reasoning is the clustering algorithm detailed above does nothing to statistically preclude clusters forming even for studies reporting different spatial effects. Consequently, the peak summary effects reported by studies, which by definition already surpass a study dependent threshold for significance, may well produce significant meta-analytic results in any incidentally formed clusters; the same is also true of meta‐regression if reported *Z* scores all correlate positively, for example, with the regressor. Consistency in reported effect size, while necessary for significance in CBRES, is not sufficient and a further step is required to prevent studies testing different hypotheses producing such incidental significant results frequently i.e. to control the family wise error rate (FWER). Furthermore, when the studies test related hypotheses such that the clusters represent truly consistent spatial effect, controlling the type 1 error rate such that the number of clusters expected under the null are quantifiably outnumbered by the significant clusters makes interpretation simple. In ClusterZ, this is performed using the FCDR.

The concept is that results declared significant by ClusterZ should be more significant than incidental results that might arise if the studies were measuring different effects. A null hypothesis based on an unbiased sample of unrelated studies with similar characteristics (subject numbers, effect sizes etc) is needed, but such samples are not readily available; indeed it is not obvious how unbiased might be defined for a sample of neuroimaging studies. In CBMA approximations to unrelated studies are computed by spatial randomisation of the reported coordinates; ClusterZ uses the algorithm detailed in section 3.1. To be specific, ClusterZ produces 4000 pseudo experiments by replacing the reported coordinates with random coordinates; while preserving the reported effect sizes, subject numbers, censoring thresholds, and covariate. For each of these, clusters are formed and inference on the effect sizes, using the likelihood ratio test, performed. If a total of *N_0_* clusters are formed by these pseudo experiments, then there are an associated set of p-values: *p_0i_* (1≤*i*≤ *N_0_*). Similarly, using the reported coordinates a set of *N* clusters are formed having an associated set of p-values: *p_j_* (sorted such that *p_1_*≤ *p_2_* ≤*p_3_*≤… *p_N_*). To control the type 1 error rate, the false cluster discovery rate is limited to a level *α* such that

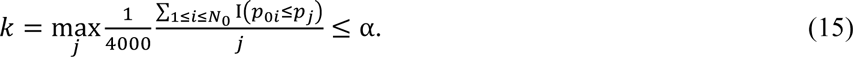

In equation (15) I(E) is an indicator function that equals one if E is true, and equals zero otherwise. The interpretation of this is that *k* is the maximum number of significant clusters such that the expected number of clusters in a pseudo experiment is at most *α*×*k*.

The FCDR is very similar to the more familiar false discovery rate (FDR) method (Benjamini 1995) on which it is based, and is exactly the same as that employed by LocalALE (Tench 2013). It imposes control such that at most a specified proportion of clusters declared significant would be expected from the pseudo experiments used as surrogate experiments involving unrelated studies. Furthermore, it controls the FWER under the null hypothesis (Tench 2013) just as FDR (Benjamini 1995).

#### 3.4.1 Family wise error rate control in ClusterZ

Beyond FCDR, it is also straight forward to control the FWE rate in ClusterZ. By taking the minimum p-value for each of the 4000 pseudo experiments, sorting them into ascending order, then picking the *α* × 4000^th^ p-value in the sorted list as the threshold for significance, the family wise error rate will be controlled at a level *α*. This is an option in the ClusterZ software, but it is conservative and has the disadvantage that there is no indicator of the proportion of clusters declared significant that are to be expected under the null, which is why FCDR is the default and recommended option.

### 4 Experiments

#### 4.1 Validation of clustering distance

To consider how sensitive ClusterZ is to Δ, numerical simulation was used. Experiments with 20 subjects were simulated with activation Z scores normally distributed with a mean of 3.79 and standard deviation 1.0; truncated to a maximum of 6.0 to avoid unrealistically large values from the tails of the normal distribution. The Z scores were truncated below 3.79 so that half of the generated coordinates were censored on average. This censoring simulates a large effect (Eickhoff 2016) to avoid false negatives as the purpose here is to demonstrate the clustering method, rather than the sensitivity of the ClusterZ. Each study included an average, after censoring, of 10 randomly (uniformly from a GM mask) distributed coordinates, and up to 5 (depending on the censoring) coordinates spatially distributed about known clustering points; this spatial distribution was Gaussian, using a standard deviation of the distances from the centre points of 4.5mm corresponding to a higher limit (least dense) of standard deviations measured in clusters detected in CBMAs (Eickhoff 2016). Experiments were generated with 20, 50 and 100 studies representing small to large meta-analyses. Clustering was performed with ϕ(Δ)=0.5, ϕ(Δ)=0.1, ϕ(Δ)=0.05, and ϕ(Δ)=0.01 for each number of studies. It was expected that for smaller clustering distances the clusters would fail to form. Clustering was also performed for fixed clustering distance, with Δ deduced such that ϕ(Δ)=0.5 for the 20 study experiment. For this fixed Δ it was expected that the clusters would grow by recruiting the random coordinates as the number of simulated studies was increased.

##### 4.1.1 Validation of clustering distance Results

Figure (1) shows the effect of the clustering distance. As expected when the overlap fraction is ϕ(Δ)=0.01 the clusters can fail to form because Δ is too small to cause coordinates to overlap, and where they do form they are fractured. For ϕ(Δ)=0.05 the clusters have formed, but again can be fractured so that the clusters are broken into fragments. For a wide range of ϕ(Δ)=(0.1 to 0.5) the clusters successfully form so, at least for this data, the method is not overly sensitive to the clustering distance. This is because the coordinates in the true clusters are generally considerably more densely packed than the between cluster coordinates.

**Figure 1.**
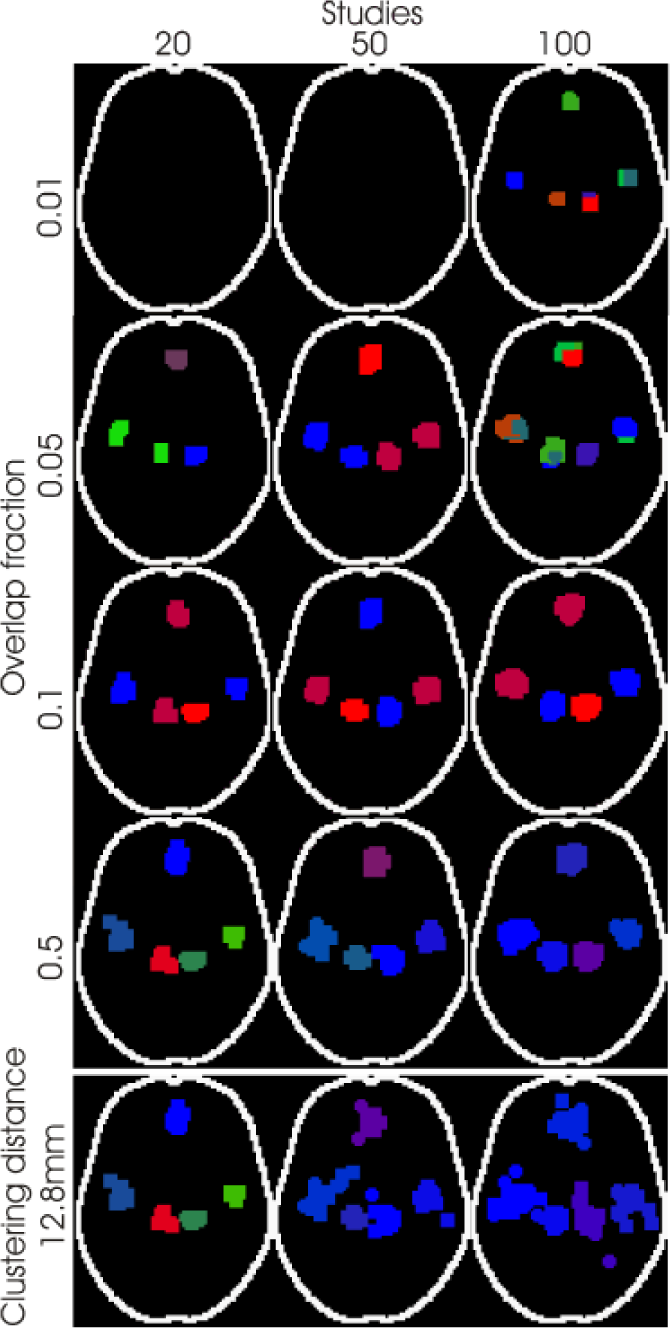
Showing the effect of clustering distance on simulated data with 5 clusters. Significant clusters are represented by cube markers for each coordinate in the cluster, and different clusters are indicated by different colour markers. For small overlap fraction (top), the clusters can fail to form. For fixed clustering distance, the clusters grow to erroneously include study specific coordinates (bottom). Importantly, allowing the clustering distance to vary with the number of studies produces similar results for large and small meta-analyses.

The important feature of the adaptive clustering scheme is that the clusters are generally independent of the number of studies in the analysis. This does not hold when the clustering distance is held fixed, as seen on the bottom row of figure (1). As expected fixing the clustering distance recruits the between‐cluster coordinates into the true clusters as the density increases with the number of studies, leading to a paradoxically increasing number of false positives for larger meta-analyses. It is important to note that this is not a feature specific to ClusterZ, as it will affect any method employing a fixed FWHM; the clustering distance equivalent. Indeed this has previously been shown to happen in the ALE algorithm (Tench 2014, Eickhoff 2016).

#### 4.2 Estimation

The necessary step in performing CBRES meta-analysis is to estimate the effects, only then can inference using the likelihood ratio test be valid. To demonstrate the utility of equations (7-12) numerical simulation was performed. Samples representative of effects in a single cluster were generated using the model

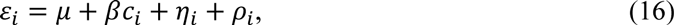

where the within-study error is

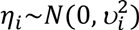

and the between-study error is

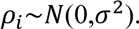

Samples of parameters *μ* and *β* were selected at random (independently) from uniform distributions with range −1 to #x002B;1, and *σ* as selected from a standard uniform random distribution but constrained to be 0.1≤*σ*≤1. To simulate *N_s_* studies, *N_s_* samples of *η_i_* and *ρ*_i_ were generated using the Box-Muller algorithm (Box 1958). For the purpose of this experiment the covariate *c_i_* was set equally spaced between −*a* and +*a*

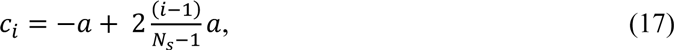

which is centred on zero and has a standard deviation of

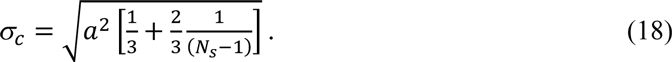

To simulate censoring any sample *ε_i_* with magnitude <*T* was removed.

A pragmatic (given knowledge of previous coordinate based meta-analyses) set of model values was employed: *n^*^*=20, *N_s_*=20, and censoring threshold *T*=3.79/√n* (which represents a commonly used uncorrected p-value threshold of 0.0001). The covariate range (−*a* to *a*) was set such that *σ_c_*=1, which according to the SLR model makes the standard errors of parameters *μ* and *β* equal. The model was generated 100 times and the estimated parameters plotted against the true parameters. It was expected that the number of studies improves the estimates, therefore a further similar experiment with *N_s_*=100 studies, representing a large MA, was simulated and results plotted.

##### 4.2.1 Estimation Results

Plots of grand mean, between-study standard deviation, and regression coefficient against their maximum likelihood estimates are shown in figure (2). MLE has been successful, over a pertinent range of parameter values, in the presence of censoring typical of whole-brain neuroimaging studies. The standard errors on the estimates are, as expected, smaller for 100 studies than for just 20.

**Figure 2.**
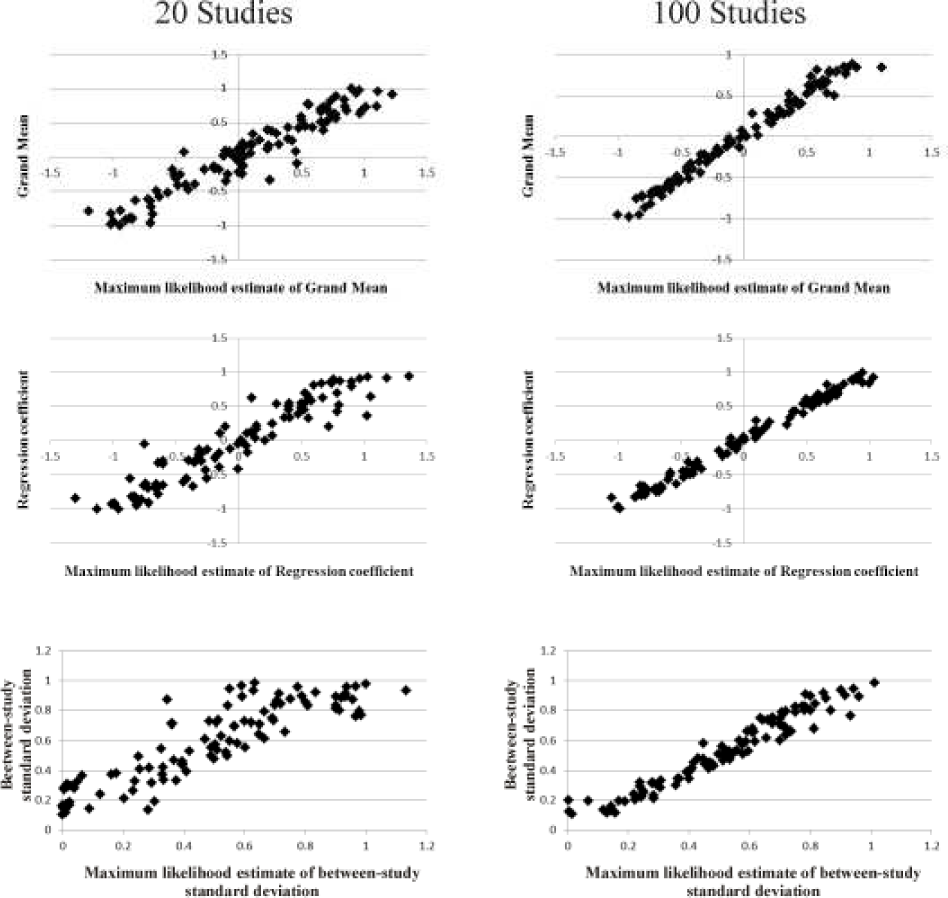
Utility of MLE to estimate the model parameters in the presence of censoring at *Z*>3.79(p≤0.0001). Estimates, as expected, are more accurate for many studies.

#### 4.3 Type 1 error control

To confirm that FCDR controls the FWER under the null hypothesis, pseudo experiments were generated by randomly placing coordinates, independently and with uniform probability, within a GM mask. This simulation used 40 studies, and the *Z* scores were sampled at random from a Gaussian distribution with mean 3.79, standard deviation 1.0, and truncated such that 3.79≤∣*Z*∣≤6; this simulates *Z* scores from studies reporting significant and censored activation (for example). Therefore, incidental clusters formed by the random coordinates may be expected to have a significant positive mean, yet there should be few significant results with FWER control. Five hundred such experiments were analysed and the number that produced significant results counted while controlling the FCDR at 0.01, 0.05, and 0.1 representing conservative, typical, and liberal settings respectively. It was expected that the pseudo experiments would produce significant results around 1%, 5%, and 10%, respectively, of the time.

Control of the FWER is important, but does not then place a quantifiable limit on the number of clusters, from those declared significant, that might be expected under the null hypothesis. This is an aim of FCDR. To test this, experiments involving known numbers of true clusters (1, 3, and 5) were simulated; with 500 simulations per experiment. Since the point of the experiment is control of false positive results, forty studies were simulated, providing sufficient statistical power to avoid many false negative results. As above, all *Z* scores were sampled at random from a Gaussian distribution with mean 3.79, standard deviation 1.0, and truncated such that 3.79≤∣*Z*∣≤6. Each study had twenty random coordinates, which was reduced to 10 on average after censoring. Added to these were a set of true clusters distributed about fixed Talairach (Talairach 1988) coordinates; the spatial distribution was Gaussian, using a standard deviation of the distances from the centre points of 4·5mm corresponding to a higher limit (least dense) of standard deviations measured in clusters detected in CBMAs (Eickhoff 2016).

To compare with the ALE and ES-SDM algorithms, 50 experiments were generated (as detailed above) for each of 0, 1, 3, and 5 fixed clusters and saved in the ALE and ES-SDM formats; these files are included as supplemental material. The distributions of clusters detected using ALE and ES-SDM were plotted as histograms, along with equivalent results from ClusterZ using the experiments detailed above with FCDR 0·05. FDR of 0·05 was used with a minimum 200mm^3^
cluster size for ALE (Eickhoff 2016), and the recommended uncorrected p-value 0·005 and minimum cluster size 10 voxels was used with ES-SDM (Radua 2012). Ideally the recommended cluster based threshold method would have been employed with the ALE algorithm, but the execution time (around one day per experiment) prevented its use for the 200 experiments performed. Nevertheless, the limitations of FDR in the context of CBMA are well understood (Eickhoff 2016) and can be considered in the comparison.

##### 4.3.1 Type 1 error control Results

Five hundred pseudo experiments, with zero fixed clusters, were processed. A histogram of the number of clusters detected are shown in figure (3). At FCDR of 0.01 the total number of experiments declaring significant clusters was 9/500, which is an estimated FWER of 0.018. Similarly at FCDR of 0.05 and 0.1 the total number of experiments declaring significant clusters was 24/500 and 54/500 respectively, representing family wise error rates of 0.048 and 0.11. This experiment suggests that FCDR is able to control the FWER under the null hypothesis, just as the FDR scheme it is based on. The ability of FCDR to control the type 1 error rate was also tested in experiments with up to 5 true significant clusters, and also shown in figure (3). The number of clusters declared significant was correct in at least 80% of experiments, and as expected the number of false negatives was highest for FDCR of 0.01, the number of false positives was highest for 0.1, while the typical setting of 0.05 fell between the two. This experiment demonstrates that, at least for the simulated data, FCDR does control both the FWE rate and the cluster-wise type 1 error rate.

**Figure 3.**
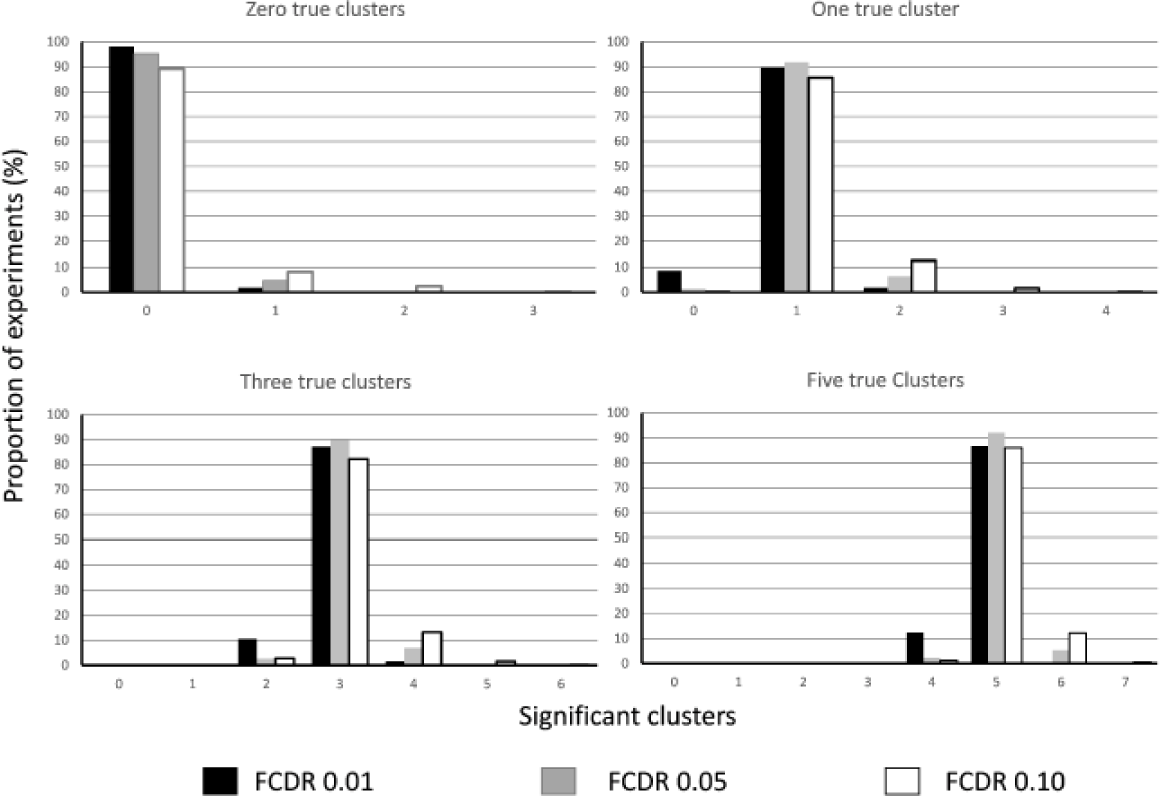
The number of clusters declared significant by ClusterZ for known numbers of clusters (0, 1, 3, and 5) and FCDR (0.01, 0.05, and 0.1).

The results of the comparison between ClusterZ and the ALE and ES-SDM algorithms is shown in figures (4) & (5). The ALE algorithm controls the FWER when there are zero true clusters since FDR was used, but the rate of false positive clusters begins to increase for 3 and 5 clusters. However, these extra clusters were small and are a known consequence of voxel-wise FDR in ALE. Taking into account the limitations of voxel-wise FDR, the ALE algorithm performs similarly with ClusterZ for this data, as can be seen in figure (5), where the small false clusters are highlighted; note the smoothness of the ALE algorithm is due to the Gaussian kernel used by the method, while ClusterZ depicts significant clusters by block markers for each coordinate contributing to the cluster. The ES-SDM algorithm, on the other hand, has been unable to demonstrate control of either the FWE rate, or the number of false clusters. This is a result of the uncorrected p-value threshold employed. Figure (5) shows the extent of the issue, with multiple clusters detected under the null hypothesis (zero true clusters) and results that are quite different to both ALE (despite the same coordinates being used) and ClusterZ, and also not constrained to the regions where the true clusters are placed.

**Figure 4.**
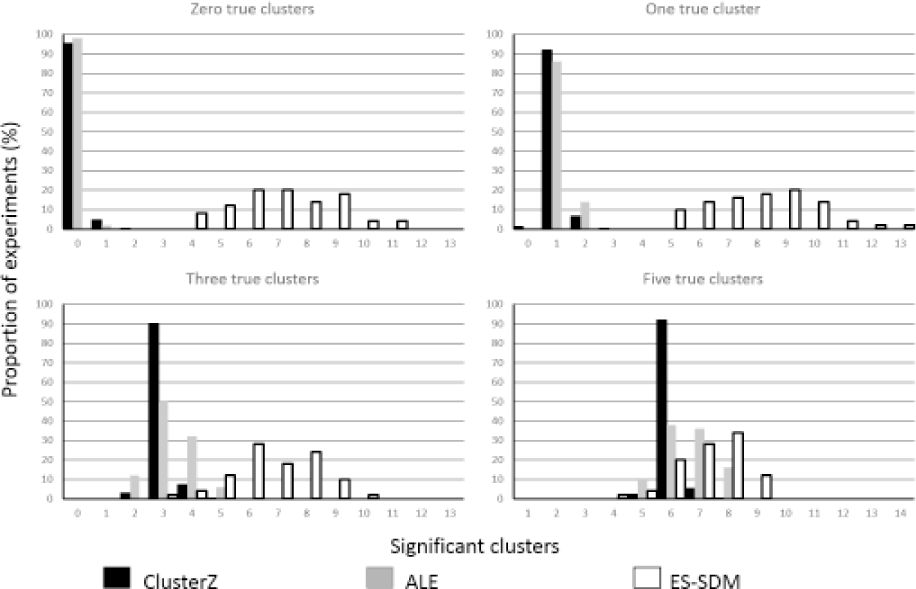
Comparison of ClusterZ, ALE, and ES-SDM using simulated data with known numbers of clusters. For ClusterZ an FCDR of 0.05 was employed, and FDR of 0.05 with a minimum cluster size of 200mm^3^was used for the ALE algorithm, and the default p<0.005 and cluster extent of 10 voxels used for ES-SDM.

**Figure 5.**
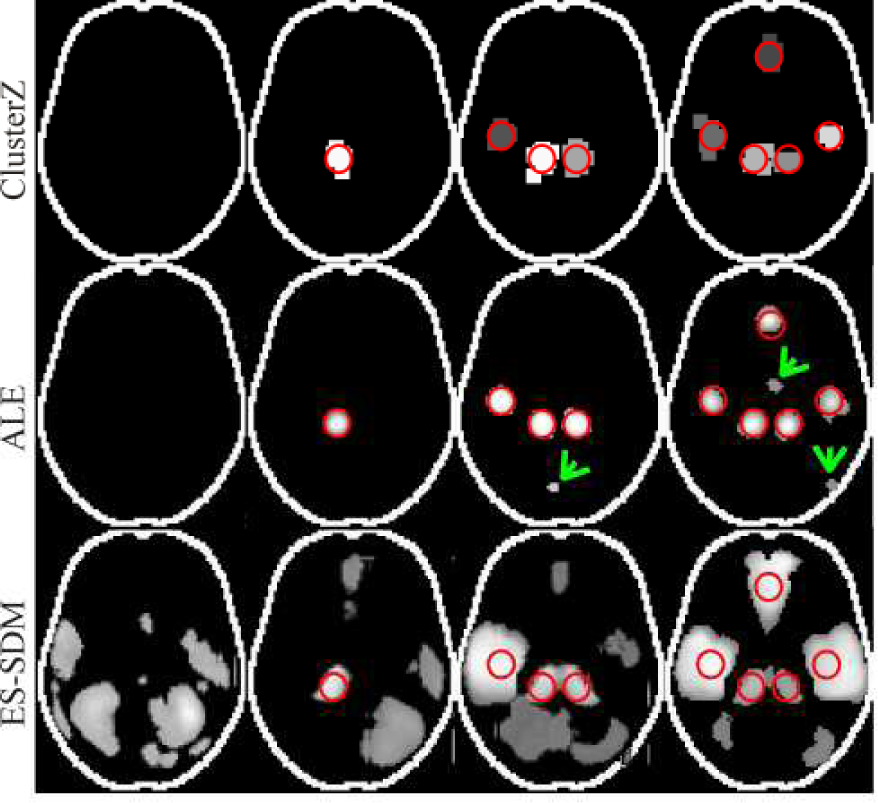
Typical results of numerical experiments with fixed numbers of clusters superimposed onto an axial outline of the brain. The red circles indicate the placement of coordinates belonging to the fixed clusters. The resulting significant clusters are shown as maximum intensity projections. For the ALE results, the arrows indicate small false clusters due to the use of voxel-wise FDR.

#### 4.4 Coordinate Based Random Effect analysis of real data

Numerical experiments verify the functionality of the algorithms and demonstrate their features, but it remains to be shown that ClusterZ performs on real data. Full meta-analyses are a study in themselves and beyond this demonstration, so data adapted from two previously published analyses (Lansley 2013, Tanasescu 2016) are used. The first is a meta-analysis of VBM studies of clinically isolated syndrome (CIS) and multiple sclerosis (MS); subjects diagnosed with CIS are at risk of developing clinically definite MS, which is known to result in grey and white matter atrophy. The studies compared grey matter density or volume of patients to healthy controls. In total there are 21 studies reporting 29 experiments comparing patients to healthy controls, and of these 4 were CIS, 16 RRMS, 3 BMS, 2 SPMS, 2 PPMS, 1 classical MS, and 1 cortical MS; details are in supplement 1. The second is a meta‐analysis of fMRI studies of mechanically induced pain in healthy volunteers. These studies compared functional activation under painful stimulus to activation at rest or innocuous stimulus. In total this includes 24 studies involving 318 volunteers; details are in supplement 1. Each analysis was performed using ClusterZ (FCDR 0.05 FWE 0.05), ALE (default cluster forming threshold p<0.001, cluster‐ level FWE 0.05, 1000 iterations), and ES-SDM (default uncorrected p<0.005, minimum cluster size 10 voxels).

##### 4.4.1 Real data CBRES Results

ClusterZ detected 6 significant clusters in the MS data using FCDR, which are listed in table 1; automatic Talairach labelling was performed as detailed in (Lancaster 2000) to locate the anatomy implicated by the clusters, but most covered more than one Talairach structure as reflected in the table. Using ClusterZ FWE method reduced the significant clusters to 3, but it is worth noting that they are identical to the respective three from the FCDR analysis; clusters, formed by the clustering algorithm, are independent of the threshold level in ClusterZ unlike voxel-wise analyses. The ALE algorithm has declared 5 significant clusters in in the MS data, which are very similarly located to those detected by ClusterZ. One cluster (left putamen) was detected by ALE but not ClusterZ at FCDR 0.05. However, a useful feature of ClusterZ is that the FCDR is estimated for each cluster detected, allowing post-hoc analysis of clusters marginally beyond the threshold. In this instance, the putamen structure was detected by ClusterZ at an FCDR of 0.06. The similarity between the ALE and ClusterZ results extends somewhat to the characteristic cluster shapes, as can be seen by comparing rows a & b with row c in both figure (6) and figure (7), however the ALE clusters tend to be smaller. This is a result of the different way that ALE represents clusters (voxel-wise activation likelihood measure) compared to ClusterZ, which, in the absence of a continuous density based measure, depicts the coordinates contributing to the cluster as a cube marker (with the colours indicating distinct clusters). Comparison with the ES-SDM algorithm is not straight forward. ES-SDM declares clusters that are comparatively large, and having different centroids; this is evident from both figure (6) and figure (7). Where possible the reported cluster centroids were matched to those detected by ClusterZ and ALE and reported in table 1.

**Figure 6.**
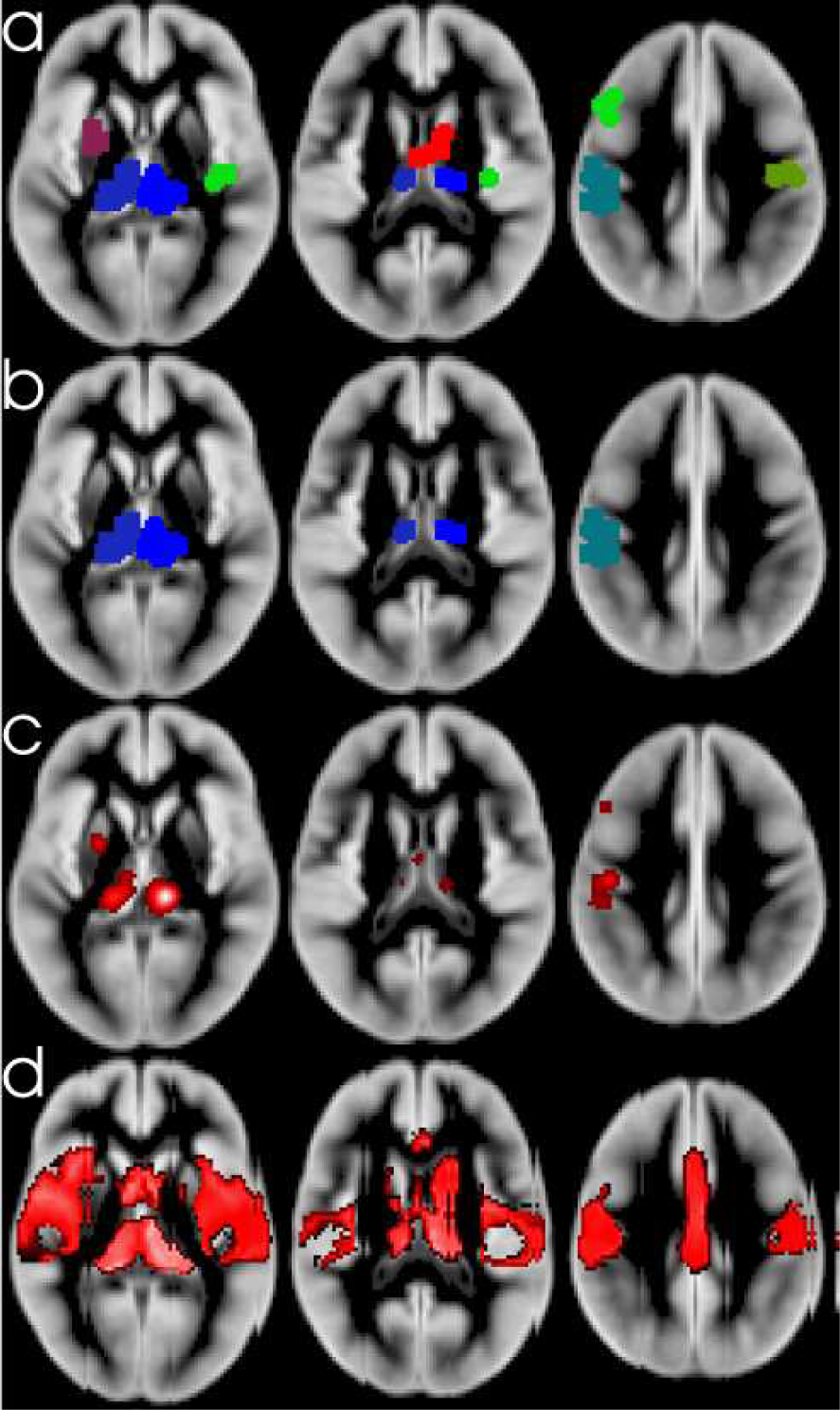
Significant clusters detected in the MS coordinates. a) is ClusterZ using FCDR 0.05, b) is ClusterZ using FWE 0.05, c) is ALE algorithm employing p<0.001 cluster forming threshold and cluster threshold of 0.05 (FWE corrected), and d) is ES-SDM using the recommended p<0.005 threshold and cluster extent of 10 voxels.

**Figure 7.**
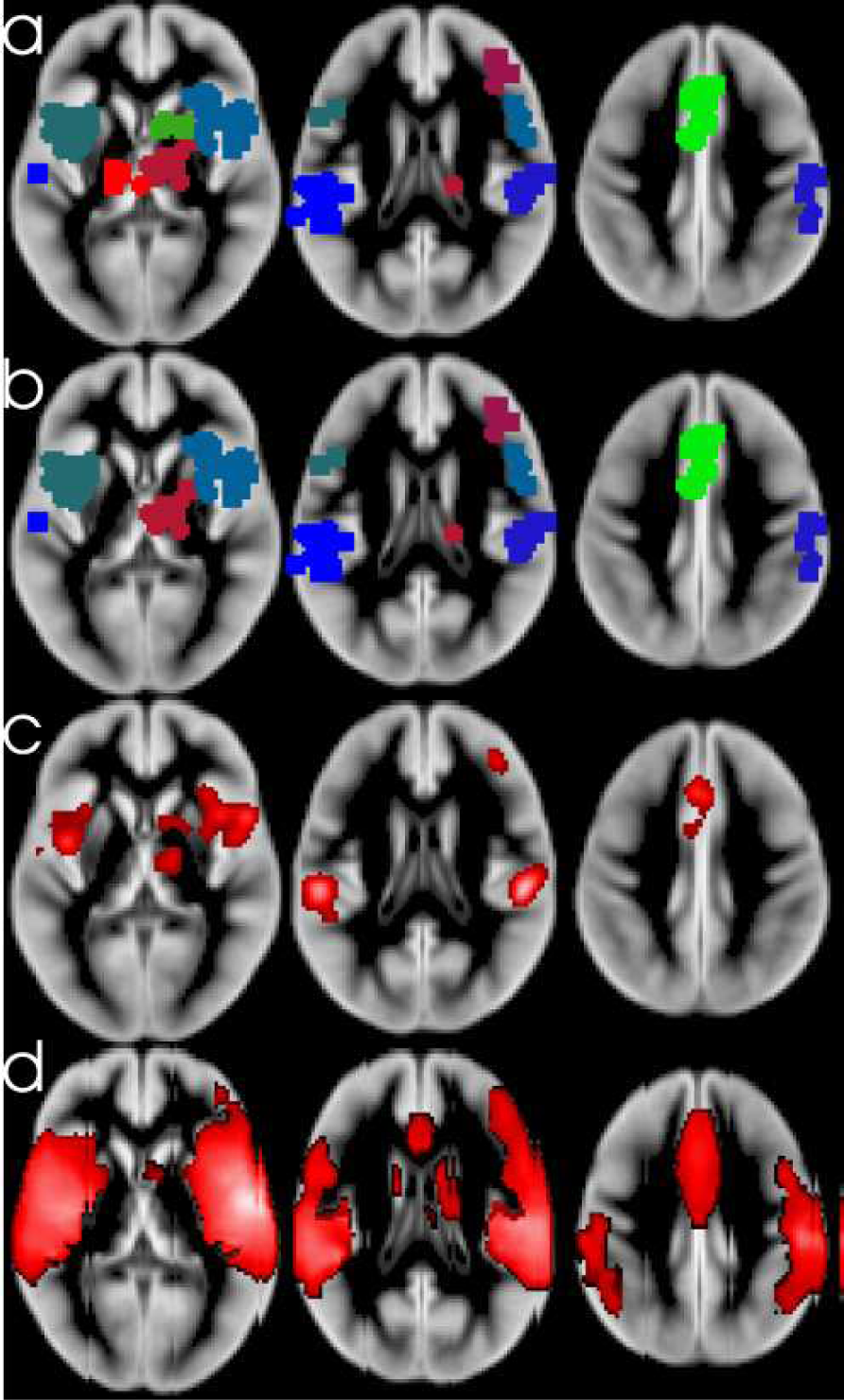
Significant clusters detected in the pain coordinates. a) is ClusterZ using FCDR 0.05, b) is ClusterZ using FWE 0.05, c) is ALE algorithm employing p<0.001 cluster forming threshold and cluster threshold of 0.05 (FWE corrected), and d) is ES-SDM using the recommended p<0.005 threshold and cluster extent of 10 voxels.

**Table 1.** Significant clusters with their estimated grand mean effect size and between-study standard deviation, and their centre Talairach location, for the MS data. The results are shown to compare ClusterZ, ALE, and ES-SDM methods. Clusters marginally beyond significant indicated by * (ClusterZ only).

| Talairach | Grand mean<br>($\mu$), between<br>study $\sigma$ | ClusterZ FCDR | ClusterZ FWE | ALE | ES-SDM |
| --- | --- | --- | --- | --- | --- |
| Right Thalamus | -1.22, 0.32 | 10.6 -26.6 7.8 | 10.6 -26.6 7.8 | 10.7 -26.7 7.9 | 6 -16 13 |
| Left Thalamus | -1.18, 0.31 | -12.5 -27.4 5.4 | -12.5 -27.4 5.4 | -12 -25.4 6.7 | - |
| Left BA 3/4/40 | -0.75, 0.35 | -48.4 -21.1 40.3 | -48.4 -21.1 40.3 | -46.9 -23 40.8 | -36 -26 43 |
| Left BA 9 | -0.55, 0.35 | -44.2 14.6 32.1 | - | -44.7 14 32.3 | - |
| Right Thalamus/Caudate | -0.56, 0.48 | 5.1 -3.2 17.4 | - | - | - |
| Right BA 3/4 | -0.59, 0.42 | 44.3 -16.4 42.0 | - | - | 50 -20 50 |
| Left Putamen | *-0.55, 0.50 | *-23.8 1.4 4.6 | - | -23.5 0.7 5.6 | - |
| Right Insula | - | - | - | - | 32 -1 15 |
| Corpus Callosum | - | - | - | - | 1 16 19 |

ClusterZ declared 9 significant clusters with the pain data using FCDR; table (2) and figure (7). Post‐ hoc analysis revealed a tenth cluster just beyond significant at FCDR 0.06. This is reduced to 7 clusters when FWE is employed. ALE agreed on 8 of these. Again the ES-SDM results were somewhat different, and matching cluster centres in table 2 was not possible for most. Differences between ClusterZ and ALE, on the one hand, and ES-SDM on the other, may relate to the large (20mm) FWHM used by ES‐ SDM (~10mm is used in ALE) and the fact that a liberal 0.005 p-value threshold is employed.

**Table 2.**
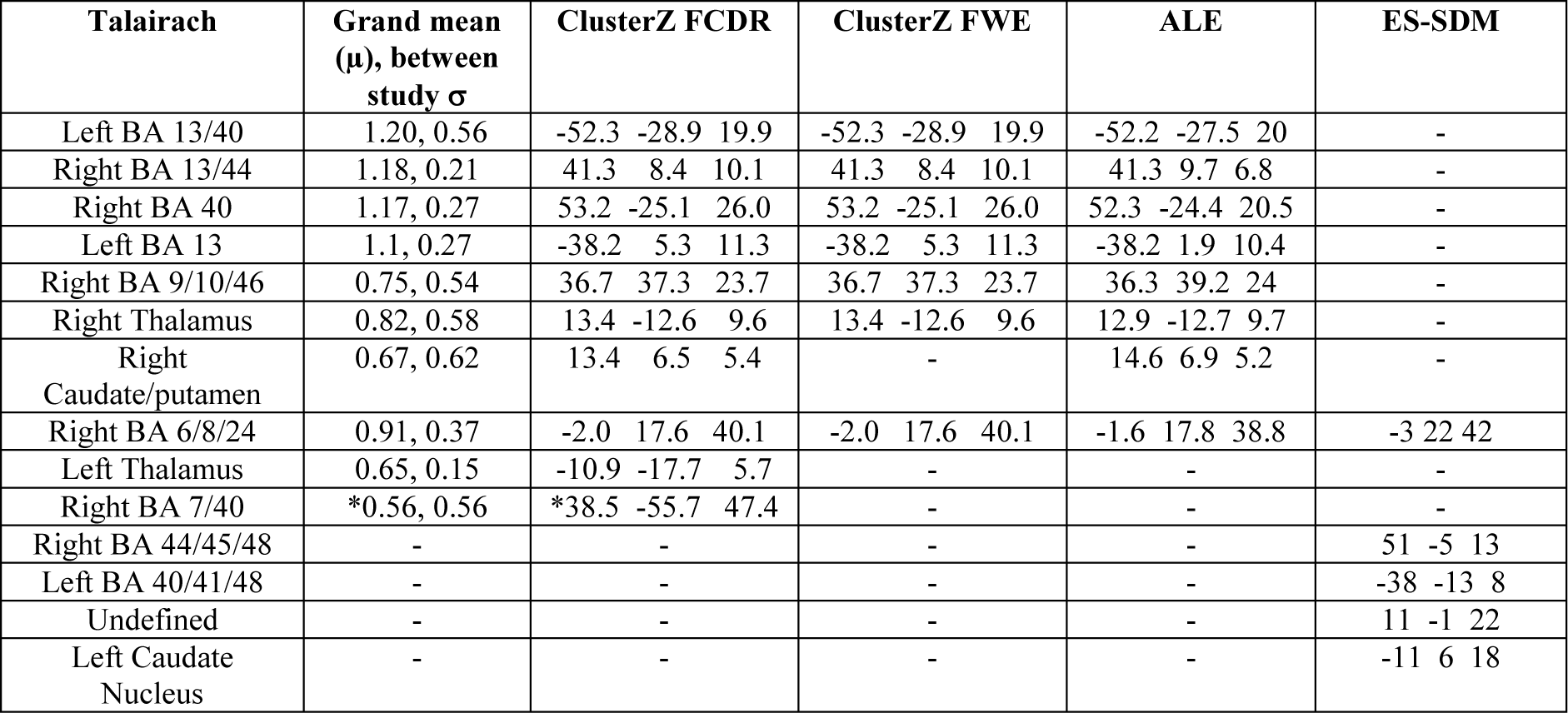
Significant clusters with their estimated grand mean effect size and between-study standard deviation, and their centre Talairach location, for the pain data. The results are shown to compare ClusterZ, ALE, and ES-SDM methods. Clusters marginally beyond significant indicated by * (ClusterZ only).

An important aspect of MA is data checking and forest plots are the standard way to visualise the data in context, which is invaluable for quickly identifying problems. Data checking is particularly important for ClusterZ, where multiple meta-analyses (one per cluster formed) are performed. The use of random effects meta-analysis in clusters means that forest plots become viable, and code to draw the plots is automatically generated for use with the R statistical package (Team 2008). Figure (8) shows forest plots from the most significant clusters from the MS and pain analyses. These indicate the reported effect sizes and a range deduced from the within-study standard deviation, an overall estimate of grand mean and standard deviation, and the p-value for the cluster. Censored studies are also depicted in the plots by empty circle markers and dashed line range indicators derived from the study thresholds. For these studies the markers do not indicate the contribution to the estimate of the model parameters, but the dashed lines indicate the range over which the contribution is computed using equations (9-11).

**Figure 8.**
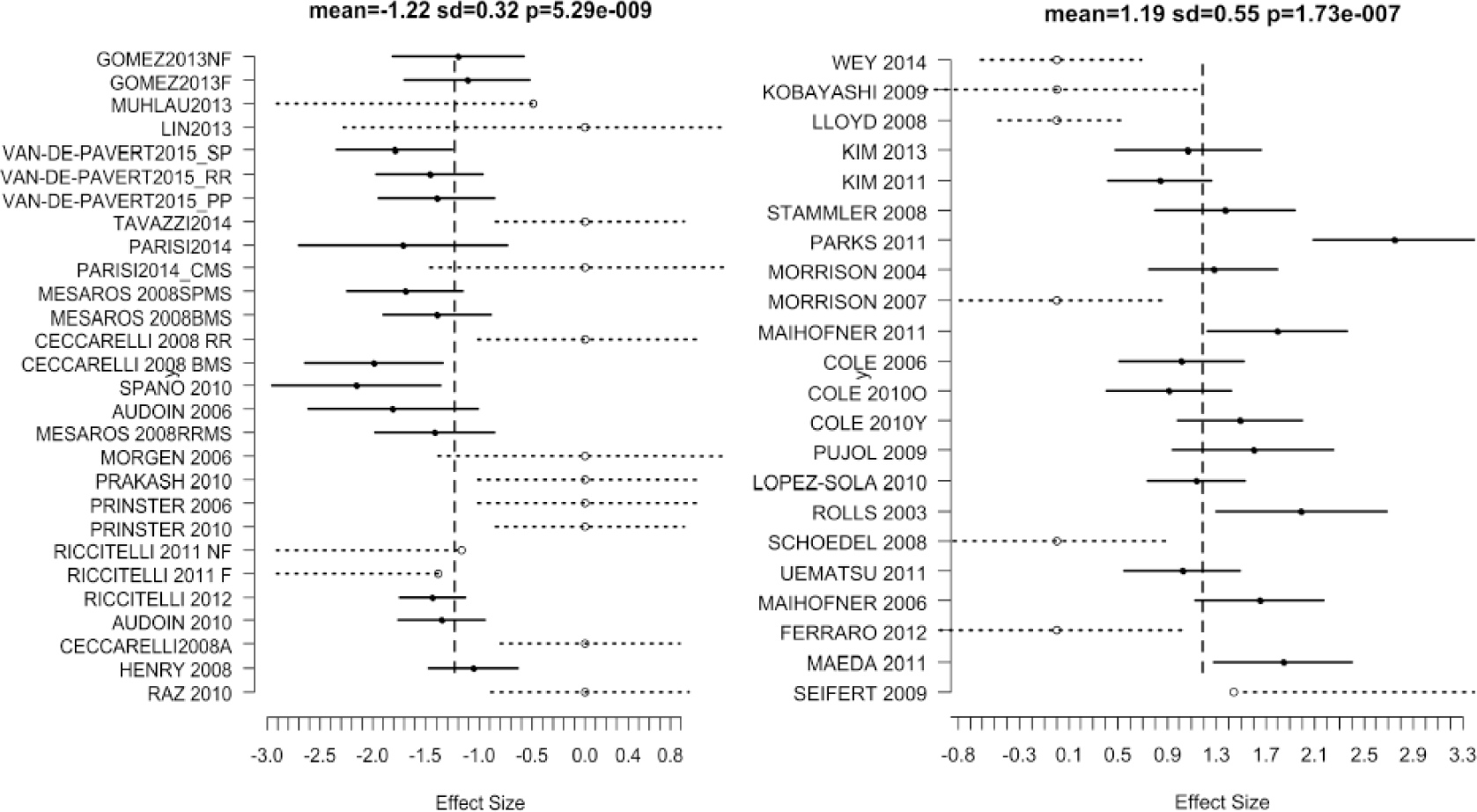
Forest plots of the effect sizes in the most significant cluster from the MS (left) and pain (right) meta-analyses. Solid circle markers indicate the effect size reported by the study in the cluster, while the confidence intervals are depicted as solid horizontal lines spanning ±1.96 times the within study standard deviation of the effect size. Censored values are indicated by open circle markers and the intervals by dashed lines (‐‐‐o‐‐‐); these are determined by the study thresholds and indicate regions where the likelihood contributions are computed using equations 9, 10, or 11.

### 5 Discussion

Here a method of performing coordinate based random effect size meta-analysis and meta-regression was presented. ClusterZ utilises the *t* statistic (or *Z* scores) and coordinates, typically reported by functional MRI or VBM studies, to detect where studies report effects consistently. The reported statistics are not suitable effect measures for meta-analysis directly, but they can be transformed (equations 1-5) into an approximate effect size that is suitable for a random effects meta-analysis. The advantages of ClusterZ include estimates of effect size, the possibility of meta-regression, and that cluster-wise statistical significance is determined by the reported effect rather than the density of clustering, which is not so interpretable. Furthermore, analysis at the cluster-, rather than voxel-, level makes type 1 error control using the easy to interpret false cluster discovery rate feasible.

ClusterZ relies on multiple established methods to perform its analysis. Firstly, clusters are formed based on the DBSCAN algorithm, which aims to differentiate true clusters from study specific effects. The clustering algorithm has a parameter that is analogous to the FWHM parameter in CBMA algorithms, but in ClusterZ this automatically adapts to the experiment to avoid false negatives when there are few studies and avoid false positives when there are many studies; the fixed FWHM used in other algorithms paradoxically increases the false positives for increasing numbers of studies. Estimation of model parameters is based on the standard statistical technique of maximum likelihood estimation, which also allows for censoring. The subsequent inference on the parameters is based on the generalised likelihood ratio test, which has well known statistical properties in the asymptotic limit. Finally, principled control of the false positives clusters is based on the popular FDR method, which controls the FWER under the null hypothesis, and then limits the proportion of clusters expected under the null hypothesis to a user specified level; by comparison the ES-SDM method employs no disciplined control of the false positive clusters. This is perhaps the most important feature of ClusterZ, since false positives results might propagate and even amplify the very issues that motivated the meta-analysis in the first place.

The adaptive clustering algorithm was tested on simulated data having known numbers of true clusters. Properties of true clusters have been empirically established (Eickhoff 2016), which was used to define a simulated true cluster with coordinates having a standard deviation of 4.5mm about a centre point. The successful formation of the true clusters depends on the clustering distance parameter Δ which is automatically estimated from the data. The results showed that the true clusters would form for a wide range of clustering distance, implying that ClusterZ does not critically depend on its exact value. Small values of clustering distance did prevent the clusters forming, or caused them to fracture, but this would be true of any other CBMA algorithm. Keeping Δ fixed was shown to increase cluster sizes as the number of studies increases indicating a lack of convergence onto a steady-state result for larger analyses. This is not specific to ClusterZ, but is a feature common to algorithms using fixed FWHM (the analogue of Δ) (Tench 2014, Eickhoff 2016). An important feature of ClusterZ, compared to other algorithms, is therefore that the results are convergent and escalating false positives are avoided for larger analyses.

Maximum likelihood estimation was used to estimate model parameters in the presence of censoring. MLE is one of multiple methods of performing meta-analysis, and has been shown to perform well in comparison (DerSimonian 1986). Numerically MLE is a nonlinear problem, with no guarantee of convergence to the global maximum. However, because ClusterZ computes estimates only at clusters, rather than voxels, the algorithm is very efficient (~1minute for typical analyses such as the MS and pain studies) so considerable computations effort is dedicated to finding the global maximum solutions. Numerical experiments confirmed its validity, being able to accurately estimate the parameters of the models represented by equation (6).

The features of FCDR were also demonstrated by numerical simulation, which showed that the number of pseudo (involving randomly generated coordinates) experiments declaring significant results is as required for control of the FWER. Having controlled the FWER the user has a measure of confidence that a significant experiment is not a false positive. Furthermore, FCDR adapts to the data to quantify a limit on the proportion of significant results that are expected under the null hypothesis. This was demonstrated on experiments containing a known number of true clusters. This is the expected behaviour of FDR, and therefore FCDR, which was designed for experiments where control of the FWER is undesirably conservative.

By comparison the ALE algorithm performed similarly with simulated data, in that it controlled the family wise error rate and correctly detected the true clusters. One limitation of the ALE experiment was the use of voxel-wise FDR, which caused multiple false positive clusters. This is a known issue, and one that is fixed with the new cluster based FWE algorithm that is now recommended (Eickhoff 2012); unfortunately the long execution time made it impracticable for this experiment. While ALE and ClusterZ performed similarly for the simulated data presented, this would not have been the case if experiments with both activations and deactivations were simulated since ALE can only process one or the other while ClusterZ can process both. The comparison with ES-SDM is not straight forward. The algorithm uses a large fixed FWHM (20mm) and a liberal uncorrected p-value to declare significant results. This resulted in lack of control of the FWE and false positive clusters. It should be noted that the coordinates used for the 200 ALE experiments were exactly the same as those used for the ES-SDM analyses, and are included as supplementary material for validation.

Analysis of real data cannot be considered validation as the results are unknown. Nevertheless, it is important to demonstrate the output of ClusterZ. Furthermore, the analysis of real data provides an opportunity to qualitatively compare ClusterZ to the popular ALE and the ES-SDM algorithms. The meta-analysis of VBM studies of MS patients revealed some of the clusters that have been demonstrated to be the most consistently atrophied in a recent study comparing software packages for volumetric analysis in relapsing remitting MS (Popescu 2016). A further example using real data analyses functional studies of healthy controls receiving painful mechanical stimulus. The ALE algorithm produced results similar to ClusterZ, but the data largely include a single effect sign (all GM loss in the MS data, and mostly activation in the pain data). If the data were a mix of activation and deactivation, ALE would necessitate two separate analyses, which statistically is a type of censoring not considered by the algorithm that could lead to bias. Comparison to the ES-SDM algorithm showed that lack of type 1 error control can produce false positive results. This is to be expected with uncorrected p-values, and has been demonstrated previously to produce apparently significant results where there are none (Tanasescu 2016).

A limitation of ClusterZ, and of the routine reporting of neuroimaging studies, is that the effect sizes (*t* statistics and Z scores) are not directly relatable to biological effect (Chen 2016). Reported grey matter volume loss in VBM, for example, would arguably be more appropriate. Prospective studies could be used to measure a more appropriate effect in a-priori regions, but first the regions need to be identified. ClusterZ, and other CBMA algorithms can do this. However, location of the regions is only one aspect of prospective study design as a sample size calculation is a requisite. ClusterZ estimates a distribution of effect sizes for each of the regions found to be consistently reported across studies, which might then be used for power calculations (Maumet 2016).

## 6 Summary

ClusterZ is a new algorithm for performing coordinate based random effect size meta-analysis, or meta‐regression, on summary results routinely reported in functional MRI studies and VBM studies. It employs disciplined control of the type 1 error rate, which is most important for a meta-analysis where rigorous statistics is necessary to gauge the strength of evidence for an effect. Advantages over coordinate based meta-analysis are that statistical significance is determined by standardised effects rather than the density of clusters, the censoring present in published summary results can be considered, and meta-regression is feasible. The effect size employed is not ideal and is limited due to the current reporting standards. Nevertheless, ClusterZ detects where the reported summary results are consistent both in terms of statistical effect size and spatial agreement, indicating target regions of interest for prospective studies of biological effect whilst also providing effect sizes that can be used to power the study.

